# Nanopore callers for epigenetics from limited supervised data

**DOI:** 10.1101/2021.06.17.448800

**Authors:** Brian Yao, Chloe Hsu, Gal Goldner, Yael Michaeli, Yuval Ebenstein, Jennifer Listgarten

## Abstract

Nanopore sequencing platforms combined with supervised machine learning (ML) have been effective at detecting base modifications in DNA such as 5mC and 6mA. These ML-based nanopore callers have typically been trained on data that span all modifications on all possible DNA *k*-mer backgrounds—a *complete* training dataset. However, as nanopore technology is pushed to more and more epigenetic modifications, such complete training data will not be feasible to obtain. Nanopore calling has historically been performed with Hidden Markov Models (HMMs) that cannot make successful calls for *k*-mer contexts not seen during training because of their independent emission distributions. However, deep neural networks (DNNs), which share parameters across contexts, are increasingly being used as callers, often outperforming their HMM cousins. It stands to reason that a DNN approach should be able to better generalize to unseen *k*-mer contexts. Indeed, herein we demonstrate that a common DNN approach (DeepSignal) outperforms a common HMM approach (Nanopolish) in the incomplete data setting. Furthermore, we propose a novel hybrid HMM-DNN approach, Amortized-HMM, that outperforms both the pure HMM and DNN approaches on 5mC calling when the training data are incomplete. Such an approach is expected to be useful for calling 5hmC and combinations of cytosine modifications, where complete training data are not likely to be available.

---

Nanopore sequencing is a third-generation technology for sequencing DNA and RNA that provides advantages over other technologies, such as its small size, long read lengths, and real-time, mobile sequencing capabilities [1, 2, 3]. Additionally nanopores are increasingly being used to detect epigenetic modifications to DNA, particularly DNA methylation [4]. The nanopore device works by running an ionic current through nanometer-wide pores. As a DNA molecule passes through the pore, the current across the pore changes in a manner that is characteristic of the molecules in the pore, namely the RNA/DNA sequence and its modifications. From measuring the current from known sequences and modifications, one can build up a supervised training dataset suitable for machine learning (ML) methods that are then able to transform future, unlabelled current signals to their corresponding sequence of bases and modifications [5].

Early studies demonstrated that nanopore sequencing could be used for the detection of epigenetic modifications in DNA by leveraging distinct current levels produced when a modified base is in the pore [6, 7]. These successes sparked the development of supervised machine learning methods for methylation calling on nanopore data [8, 9, 10, 11]. The first methylation modification tackled by nanopore technology was 5-methylcytosine (5mC), a well-studied modification due to its abundance in the human genome [12] and its links to a number of key biological processes such as aging and cancer [13, 14]. More recently there have been efforts to tackle a different cytosine modification, 5-hydroxymethylcytosine (5hmC), which is common in mammalian brain tissue, accounting for 40% of modified cytosine in the central nervous system [15]. 5hmC content in brain cells increases with age, suggesting that it is linked to neurodevelopment [16]. Early results suggest that nanopore current may be sensitive to 5hmC [6]. However, calling 5hmC accurately in the presence of other modifications has not yet been conclusively achieved, largely because of the difficulty in obtaining sufficient labelled training data for the ML-based callers [17].

## Generalization capabilities of nanopore callers

Although supervised ML methods are developed specifically for their ability to generalize to unseen examples, the notion of generalization for nanopore sequencing is nuanced. For example, one form of generalization for base calling is from one set of current observations for one *k*-mer, to slightly different current observations for that same *k*-mer arising from stochastic noise in the system. We call this *sensor generalization*, because the generalization required is owing only to sensor noise. Another form of generalization relevant to nanopore sequencing is *k-mer generalization*, wherein an ML-based caller must make accurate calls for *k*-mers that it has never seen current observations for. Analogous notions of generalization can be made for epigenetic calling, where, in addition to *k*-mers comprised of the standard nucleotides, we also consider *modified k-mers* that include methylated bases. In this case, *k*-mer generalization refers to generalizing to the combination of both bases and modifications.

When constructing a training dataset for *base* callers, it is relatively easy to generate a *k-mer complete* dataset—one in which current observations associated with all possible *k*-mers are present. This can be achieved by taking, for example, a sample of human DNA of known sequence, amplifying the DNA and running it through the nanopore. Labels for training can be obtained using alternative sequencing platforms. Consequently, it typically suffices to require only sensor generalization for base calling.

When it comes to constructing a training dataset for a particular methylation modification, it can be more difficult to obtain a similarly comprehensive dataset. This difficulty arises from the burden of obtaining high-confidence reference labels for these modifications. In previous studies of 5mC modifications, either enzymatically methylated DNA [10, 11], or the gold standard assay of bisulfite sequencing, was used to obtain supervised labels [18, 19]. However, as we move to other modifications, such as 5hmC, achieving similarly complete training data can be more difficult still. TET-assisted bisulfite sequencing (TAB-seq) and oxidative bisulfite sequencing (oxBS) are currently the standard methods for reading 5hmC at single-base resolution [20]. However, both methods are expensive and low-throughput [21, 22]; they also require high coverage to make high-confidence 5hmC calls (particularly oxBS) [20]. Additionally, beyond these sequencing challenges, rarity of certain epigenetic modifications may also present a problem, as it may be the case that not all *k*-mers containing a given modification are represented in a specific genome, and synthesizing each one is often not feasible. As the field progresses to simultaneous calling of multiple types of epigenetic modifications, achieving a complete dataset with respect to all of the modifications will become harder still [17]. Consequently, as nanopore sequencing technology is pushed to call more and more DNA modifications, we require ML-based callers that are accurate even with limited training data. In particular, the callers will require both *k*-mer and sensor generalization.

To further illustrate these difficulties, consider that for base calling, the nucleotide alphabet is of size four: `{A, C, G, T}`, whereas for a given methylation mark that can occur only on a cytosine, we expand the alphabet to size five: `{A, C, G, T, M}`. For current pore models where *k* = 6, we go from 4^6^ = 4, 096 unique *k*-mers to 5^6^ = 15, 625. Additionally, even if the pore contains only, say, six bases at a time, ML callers may be able to make use of larger contexts, such as nine, to improve calling [23]; this exacerbates the combinatorial explosion of possible *k*-mers (5^9^ = 1, 953, 125). Similarly, simultaneously calling even two distinct modification types, such as 5mC and 5hmC, would dictate an alphabet of size six, corresponding to 6^6^ = 46, 656 unique 6-mers. Herein, we restrict ourselves to *k* = 6 and to calling a single modification (5mC), although the conclusions that emerge should be equally, if not more, applicable to larger values of *k* and to situations in which there are multiple epigenetic modifications of interest.

Next we describe the two main modelling approaches currently used for nanopore-based methylation calling, and discuss how each is able or not able to perform sensor and *k*-mer generalization. Then we propose a new approach, which is a hybrid between the two existing approaches, and demonstrate its utility in the limited training data regime. Note that in both of these existing modelling approaches, and in our own, the methylation calling assumes that base calling has already been performed.

## Hidden Markov Model nanopore callers

Simpson et al. [10] developed the widely-used Nanopolish, a Hidden Markov Model (HMM)-based approach to detecting 5mC in CpG contexts. The Nanopolish HMM assumes a different current distribution for each unique *k*-mer, including distinct distributions for modified versions of a *k*-mer. For example, a *k*-mer, `CGAACG`, that has a 5mC in the fifth position, denoted `CGAAMG`, has its own mean and variance of current distribution in Nanopolish, and `MGAAMG` in turn has its own, and so forth. That is, every possible modification on top of any DNA background—a unique modified *k*-mer—has its current distribution modelled independently. The Markov transitions in the HMM ensure a coherence of calls as the DNA sequence moves through the pore. That is, if the HMM believes the last call in the sequence being pulled through the pore was a `CAMGAT`, then the next call in the sequence should be off-set by a shift of one, `AMGATX`, for wildcard `X`. Because of the independent current distributions—called emission distributions in HMM parlance—for each modified *k*-mer, the HMM-based Nanopolish approach cannot accurately make calls for modified *k*-mers not seen in the training data.

## Deep Neural Network nanopore callers

Recently, there has been a shift to using deep neural networks (DNN) for base [24, 25, 26] and methylation calling [18, 19]. In particular, Ni et al. [18] created DeepSignal by employing a bidirectional recurrent neural network (RNN) with long short-term memory (LSTM) units to construct sequence-based features, jointly with a convolutional neural network (CNN) to process the current. Liu et al. [19] similarly used an LSTM-RNN in their DeepMod, also adding a secondary neural network to account for correlation of modifications on nearby sites. These DNN-based methods have shown improved performance over the HMM-based Nanopolish for 5mC calling [27]. Importantly, because these DNN approaches do not have parameters that are *a priori* independent for each modified *k*-mer, it stands to reason that they should perform better than HMM-based approaches in generalizing to new modified *k*-mers—that is perform better *k*-mer generalization. Although it has not previously been shown, we will demonstrate that this is indeed the case.

## A novel hybrid HMM-DNN approach to methylation calling

Although we will show that the DNN approach has better *k*-mer-generalization than the HMM approach, we hypothesized that combining the two modelling approaches may provide better *k*-mer generalization yet, and therefore better robustness to incomplete training datasets. The HMM is inherently a low capacity model, with relatively few parameters, while DNNs typically have order of magnitudes more parameters, require vast amounts of data to train, and also potentially days to weeks of architecture search to find a useful model. We hypothesized that it may be possible to get the best of both these worlds. Our approach, Amortized-HMM, first trains a Nanopolish HMM on the training data that is available, yielding an emission distribution for each modified *k*-mer in the training data. Next, we use these results to train a feedforward deep neural network (FDNN) to learn a mapping from modified *k*-mer to HMM emission distribution parameters. Finally, we use the Nanopolish HMM, with any missing modified *k*-mer emission distributions imputed by the FDNN. In this strategy, information sharing between emission distributions is possible by way of the FDNN. Consequently, we say that we are amortizing the emission distributions, hence the name, Amortized-HMM. Although one could use only the FDNN emission distribution parameters for all modified *k*-mers, this did not perform as well (see Methods and Supplementary Figure S1). In addition to developing this hybrid model, we also developed a new algorithm for choosing which *k*-mers to use for training in the *k*-mer incomplete setting.

Next we present a series of experiments comparing and contrasting our proposed hybrid approach to pure DNN and HMM approaches, across a range *k*-mer incompleteness settings. We show that for complete training data, the DNN is best, but that as training data becomes incomplete, that our hybrid approach dominates in performance.

## Results

We focused our empirical investigation on the problem of 5mC calling, for which several high quality datasets exist, and for which existing callers have been developed with the intent of having approximately *k*-mer complete training data. This investigation serves as a proof-of-principle for harder tasks such as 5hmC calling, or joint calling of 5mC and 5hmC, and so forth, for which we could not conduct such experiments owing to the lack of complete training data motivating this very work. Our comparisons used existing datasets from naturally occurring 5mC modifications in human genomes, as in Ni et al. [18] and Liu et al. [19]. In particular, we trained and evaluated our models using two primary nanopore datasets obtained from sequencing two different human genomes, HX1 [8] and NA12878 [28]. Additionally, we obtained gold standard bisulfite 5mC labels for NA12878 from ENCODE (ENCFF835NTC) [29] and for HX1 by processing bisulfite sequencing data from the NCBI Sequence Read Archive (PRJNA301527) [8] using Bismark [30].

### Construction of incomplete training data sets

From these datasets, we constructed a range of *k*-mer incomplete training datasets in order to assess how different callers performed in different settings of incompleteness. These incomplete data sets were created within 6-fold cross-validation. By default, each training fold contained *k*-mer complete data. Next, to create a, say, 10%-complete training dataset, we compute the number of modified *k*-mers that this corresponded to, say 250 *k*-mers. In principle, we could then have simply chosen 250 of the training modified *k*-mers at random for our 10%-complete dataset. However, this would not correspond to a real physical situation owing to the fact that a single methylated site in a genome corresponds to six modified *k*-mers (for *k* = 6), all shifted from each other by one position—a structure that random sampling would not capture. To account for this physical reality, we used a slight modification of the random modified *k*-mers selection scheme, whereby we enforce that all six modified *k*-mers for that one modified site are simultaneously included in the training data. This was achieved algorithmically using an integer linear program that also accounted for the frequency in the training data of the selected samples, enabling more commonly occurring data to be better represented if desired. This makes sense when, for example, training on human genomes for ultimate deployment on human genomes (*i. e*., when the train and test distributions of modified *k*-mers are the same). However, in the case of synthesized sequences, this could be turned off. Note that each individual modified *k*-mer will generally appear many times in the training data (with distinct sensor readings), but the total number of unique *k*-mers is limited, dictated by the specified level of incompleteness. Note that test sets remained the same for every level of incompleteness and were *k*-mer complete, although in our results we decompose the accuracy among those seen in the training data and not.

We denote different levels of *k*-mer completeness by *p*. That is, *p* is the percentage of all possible unique modified *k*-mers that are present in the training data. A *k*-mer complete dataset has *p* = 100, while increasingly less complete data has increasingly smaller values of *p*. The smallest *p* we considered is five, which corresponds to fewer than 150 unique modified *k*-mers in the training data. The 20 values of *p* used can be seen on the horizontal axis in Figure 1.

**Figure 1:**
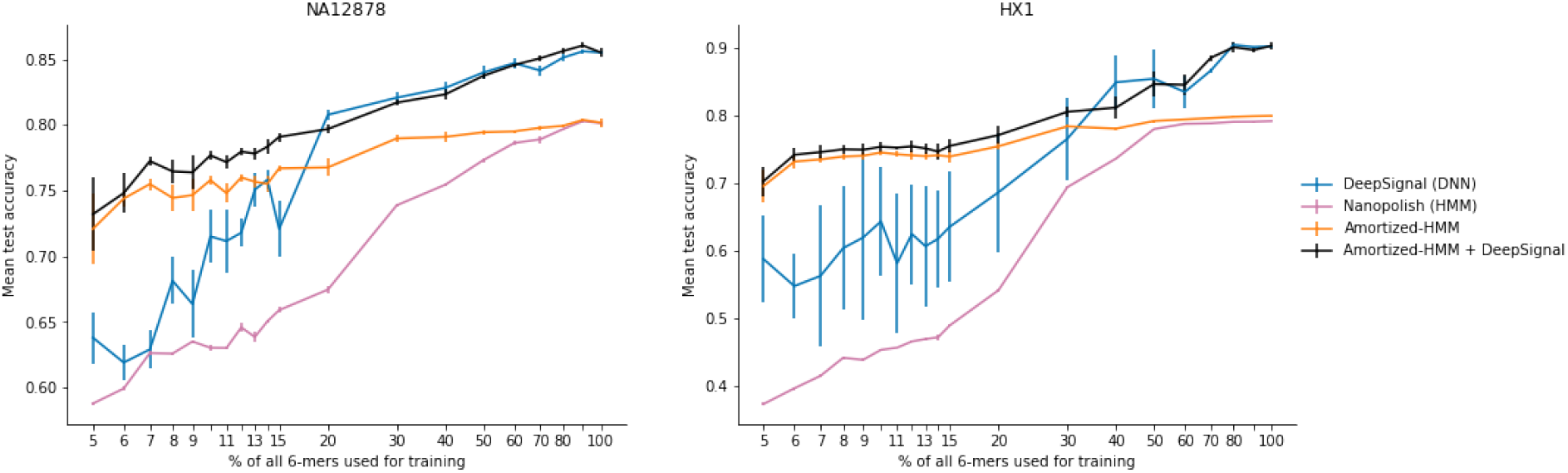
Performance of 5mC calling across different *k*-mer incompleteness regimes. Results averaged over 6-fold cross-validation, with length of error bars equal to one standard deviation across the folds. The *k*-mer complete case (*p* = 100), contains 2, 669 unique modified *k*-mers, whereas *p* = 5 contains 133. The units of test accuracy were percent modified, meaning the best accuracy was 0, and the worst was 1.

We trained three different methylation callers, described in the previous section: Nanopolish (HMM), DeepSignal (DNN), and Amortized-HMM, on each *k*-mer incomplete dataset. For the first two we used code provided by the authors. No model selection or architecture search was performed for these methods because Nanopolish does not require it, and architecture search for DeepSignal had already been performed for the *k*-mer complete setting. For our approach, Amortized-HMM, we similarly performed architecture search only in the *k*-mer complete setting.

### Accuracy of 5mC calling across a range of *k*-mer-incompleteness

On both datasets, the performances of Nanopolish and Amortized-HMM were very similar for high values of *p* (Figure 1). This is to be expected, since when *p* is close to 100, Amortized-HMM does not need to impute many emission distributions; rather it can use the Nanopolish emission distributions directly. Note that even for the *k*-mer complete setting, Amortized-HMM and Nanopolish may diverge because the latter requires a minimum number of data points for each emission distribution, and otherwise sets this distribution to the default of being unmethylated. In contrast, Amortized-HMM would use the FDNN-predicted emission distribution. Consistent with earlier results, we find that DeepSignal outperforms Nanopolish in the *k*-mer complete setting [18, 27], and similarly outperforms Amortized-HMM in that same setting, for similar reasons.

As the training data become increasingly incomplete (Figure 1), Amortized-HMM starts to systematically outperform Nanopolish because of its ability to impute missing emission probabilities corresponding to modified *k*-mers not in the training data.

Meanwhile, DeepSignal continued to hold an advantage over the other methods for *p* higher than around 20 to 30, the cross-over point for where Amortized-HMM starts to outperform DeepSignal. We hypothesize that the diminished performance of DeepSignal with increasingly *k*-mer incomplete data arises from insufficiently diverse data to train on relative to the capacity of the model. Note that both DeepSignal and Amortized-HMM had their architectures tuned only on the basis of *k*-mer complete training data, so as to make the comparison fair. However, it is possible that performing architecture search for each of the two approaches could have altered the relative performance. To investigate this hypothesis, we performed an architecture search only for DeepSignal, at *p* = 10—a regime of large incompleteness and one where Amortized-HMM substantially outperformed DeepSignal. Although the architecture search improved the performance of DeepSignal, Amortized-HMM, critically with *no hyperparameter tuning*, always outperformed DeepSignal (Supplemental Figure S1). These results suggest that the Amortized-HMM approach is more robust to changes in training data completeness, mitigating the need to redo computationally intensive architecture searches for different deployment scenarios.

Although Amortized-HMM performed best for our main regime of interest—low values of *p*—we considered whether combining Amortized-HMM with DeepSignal could yield an all-around best performer. Therefore, we made another combined approach wherein we used DeepSignal to make predictions on modified *k*-mers that occurred in the training data, and Amortized-HMM for the rest. Indeed, this combined approach yielded an overall best method (Figure 1). In the next investigations, we do not include this approach because the intent there is to understand the behaviour of the models it is composed of.

### Decomposition into sensor and *k*-mer generalization

In order to better understand the source of each approach’s errors with respect to test modified *k*-mers that appeared in the training data (sensor generalization), and not (*k*-mer generalization), we partitioned the test sets according to whether the modified *k*-mer had been seen in the training data (Figure 2). Across both primary datasets, DeepSignal was the clear winner for pure sensor generalization, while Nanpolish and Amortized-HMM performed similarly to each other, and well below DeepSignal. On the other hand, when *k*-mer generalization was required, Amortized-HMM was the consistent winner, with DeepSignal coming in second, and Nanopolish, last. These results suggest that DeepSignal is most likely overfitting to the data when it does not have access to a complete training data set, but performs well on those modified *k*-mers that it has seen at train time.

**Figure 2:**
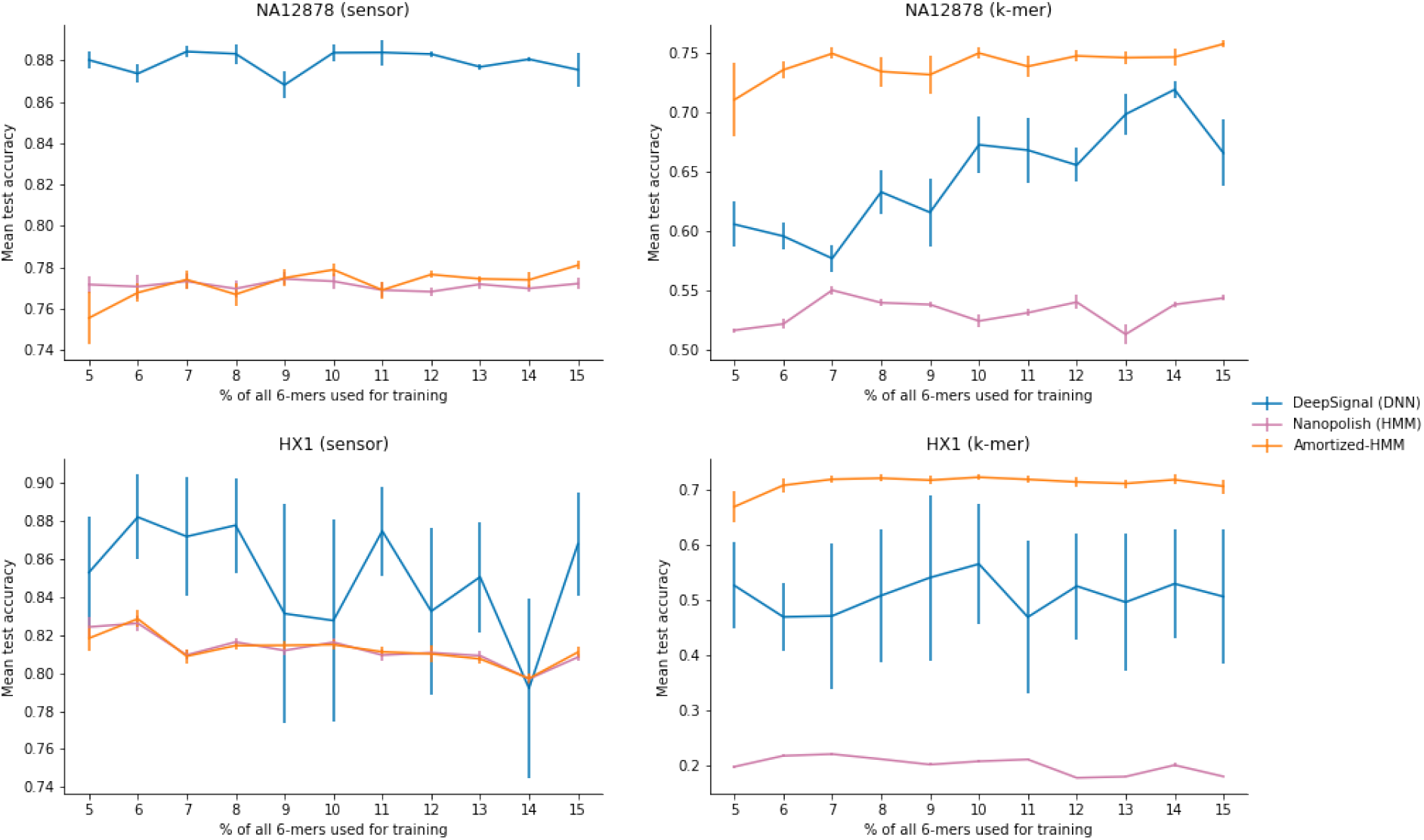
Sensor and *k*-mer generalization. Accuracy of the three approaches when the test set is broken down into modified *k*-mers observed at training time (sensor), and not observed (kmer+sensor). General figure information is the same as in Figure 1.

### Investigation of low-and high-novelty in *k*-mer generalization

In the previous section, we treated any modified *k*-mer not appearing in the training data in the same way, regardless of how similar it may have been to one of the training modified *k*-mers. However, it may be the case that this similarity plays in a role in how well the various models do. Therefore, we refer to test modified *k*-mers that were less similar to any training examples as *high-novelty* test cases, and those that are more similar (but still different) as *low-novelty*. Similarity was defined by the Hamming distance of the one-hot encoded modified *k*-mers, and similarity to the training data was obtained by averaging this quantity over all unique modified *k*-mers in the training data. Using this definition of similarity for novelty, we compared the different approaches over all values of *p* between 5-15 appearing in Figure 2, averaged over *p* (Figure 3). Although Amortized-HMM performed similarly to DeepSignal for low-novelty *k*-mers, it was far more accurate than DeepSignal for high-novelty *k*-mers. This difference in performance appears to underpin the success of Amortized-HMM over DeepSignal in *k*-mer generalization.

**Figure 3:**
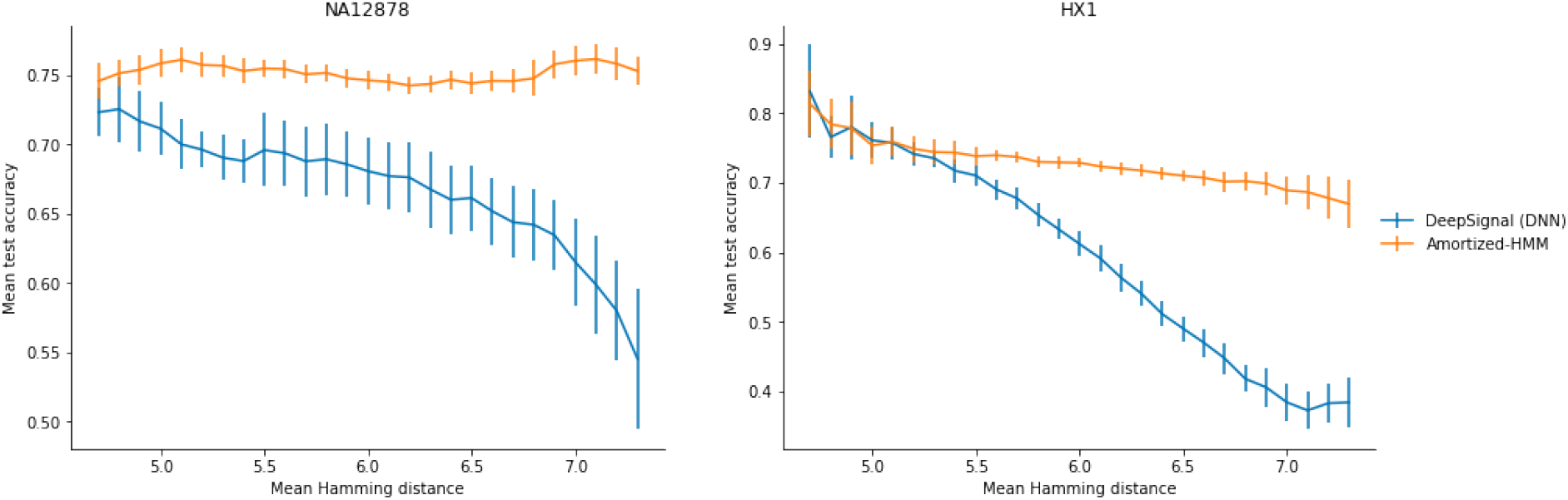
Breakdown into low-and high-novelty test set accuracy. Accuracy of DeepSignal and Amortized-HMM on *k*-mers previously unobserved during training at varying levels of *k*-mer novelty. Results are averaged over values of *p* between 5 and 15 from Figure 2). General figure information is the same as in Figure 1. Hamming distance was computed on 11-mers containing consecutive 6-mers as described in *Construction of incomplete training data sets* and *Methods*.

## Discussion

We investigated how several common modelling approaches, and our newly developed approach, for nanopore calling, are robust to generalizing to *k*-mers not seen at training time—what we refer to as the task of *k*-mer generalization. Although the DNN-based DeepSignal performed best with complete *k*-mer training data, as the training data became increasingly less and less complete, our newly proposed hybrid approach that combines HMMs and neural networks, Amortized-HMM, dominated in calling accuracy.

Although we focused our evaluation on 5mC detection, in practice, 5mC calling does not require effective *k*-mer incomplete modelling, as there already exist high *k*-mer coverage nanopore sequencing labelled datasets for 5mC. This setting was used as a proof-of-principle toward making progress on modifications for which obtaining such a dataset is not so straightforward. In particular, we are working to improve calling for 5hmC and naturally occurring combinations of cytosine modifications.

We note that both the existing callers investigated herein, and our own approach made calls having assumed that base calling has already been performed. In practice, it could be useful to combine the two tasks together for further improvements, as can be done in Guppy, the base caller developed by Oxford Nanopore Technologies [17].

## Methods

### Datasets

We trained and validated models for 5mC calling on two published Nanopore datasets. Jain et al. [28] sequenced the human genome NA12878 at 30x coverage using the Oxford Nanopore Technologies (ONT) R9.4 pore chemistry, and Liu et al. [19] sequenced the HX1 genome, also at 30x coverage and using R9.4 chemistry. In both cases, sequencing was performed on native DNA molecules containing native modifications; this is in contrast to other works where methylation was synthetically introduced in PCR-amplified samples using enzymes such as M.SssI methyltransferase [10, 11]. In addition to raw current data, these datasets included base calling results obtained from running ONT-trained base callers: Guppy v2.3.8 for the NA12878 dataset and Albacore v2.3.1 for the HX1 dataset. To obtain supervisory 5mC labels for training our caller, we followed Liu et al. [19] and Ni et al. [18]. That is, first we obtained gold standard bisulfite sequencing 5mC labels: for NA12878, these were downloaded directly from ENCODE (ENCFF835NTC) [29]; for HX1, we downloaded bisulfite sequencing results from the NCBI Sequence Read archive (PRJNA301527) [19], and pre-processed them using Bismark [30]. Then we filtered to keep only CpG sites that were (1) covered by at least 5 reads and (2) were consistently called as methylated or unmethylated across every read. This process yielded a set of CpG sites for which we could confidently assign binary (non-methylated or methylated) labels for training and evaluation purposes.

### *K*-mer selection for simulating incomplete training data

The motivation of our manuscript is that one is unlikely to have *k*-mer complete training data for many epigenetic nanopore calling problems of interest. This problem arises from, for example, limitations in experimental time and cost, or the modification of interest occurring infrequently in the genome in a given genomic context. However, since the datasets considered in this work are, by intent, *k*-mer complete, we simulated incomplete data sets by selecting only certain *k*-mers to retain. The goal of this selection procedure was to (1) mimic physical constraints of pulling a sequence through a pore (described soon), and (2) select a diverse, representative set for training so as to most clearly see the effects of incomplete training data. Note that we focused only on limiting the number of unique modified *k*-mers present in the training data, since one can assume that training data for *k*-mers with no modifications can be procured easily. For simplicity of language, we refer to the modified *k*-mers as *k*-mers in this section. Throughout all of our experiments, the number of sites in the pore at any one time, *k*, was 6, although our algorithms can be readily applied for other values of *k*. Also note that we consider only 5mC in CpG contexts, which is representative of mammalian methylation [31].

Let *S* denote the set of unique *k*-mers that contain a methylated CpG site. As described in the main text, *p* denotes the percentage of all possible modified *k*-mers that our incomplete dataset should contain. For example, if *p* = 100, then we retain all *k*-mers in *S*. For *p <* 100, requiring 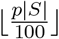 unique *k*-mers, we might consider selecting them at random. However, such an approach would yield a training set not obtainable from an actual nanopore experiment because it is not accounting for the fact that as a sequence gets pulled through the pore, the corresponding *k*-mers that arise are overlapping. Augmenting the nucleotide alphabet with `M` to represent a methylated cite, consider the 11-mer, `GATTTMGCAAC`, centered on one methylated site. This 11-mer comprises six overlapping 6-mers: in order, `GATTTM`, `ATTTMG`, `TTTMGC`, `TTMGCA`, `TMGCAA`, and `TTTMGC`, that would arise from pulling it through the pore. In our selection scheme, we therefore constrain ourselves to selecting at the 11-mer level, which directly implies a selection of six *k*-mers at once. We refer to this as a *coherent k*-mer-selection scheme, because it adheres to the physical reality of pulling a DNA strand through the pore. The random *k*-mer-selection scheme is not coherent.

To ensure a coherent *k*-mer-selection scheme, we formulate our *k*-mer selection problem as an integer linear program (ILP)—an optimization of a linear function subject to a set of linear constraints over integer variables. In our setting, the linear function takes as input the presence/absence of each 11-mer (*y*_*i*_ *∈*0, 1 for *i* = 1, …, *m*, where *m* denotes the total number of possible unique 11-mers centered on a methylated site), and returns the frequency-weighted count of the 11-mers to be retained in the data (Equation 1). The frequency weights, *w*_*i*_ *∈* [0, 1] (with 1 = ∑ _*i*_ *w*_*i*_), for any given 11-mer, are specified by the user and described below. As detailed above, the *i*^th^ 11-mer comprises a set of six *k*-mers, denoted *V*_*i*_. The presence or absence of an individual *k*-mer, *s*, is denoted by *x*_*s*_ ∈ {0, 1}. The two constraints of our ILP are that the number of selected unique *k*-mers is less than our budget, 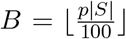 (Equation 2), and the other ensures *k*-mer-coherence (Equation 3). Altogether, our ILP is given by:

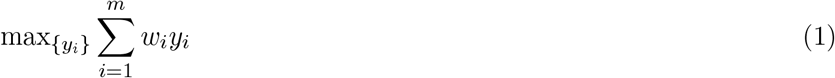

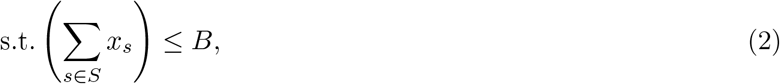

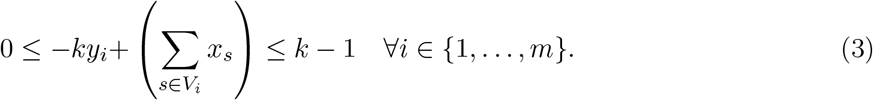

Equation 3 may be more easily understood by noting its equivalence to the *k*-way logical AND constraint, 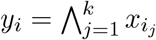, where ⋀ denotes a logical AND operator. The frequency weights could correspond to prevalence in a human genome, or could be set to uniform. In our experiments, we set the weights to be the frequencies observed in each labelled training data set. In a more realistic scenario, however, a reasonable proxy is to use the frequency of the unmodified *k*-mers; we found this also worked well (data not shown).

After solving the ILP with a standard solver (Gurobi [32]), we obtain values of *y*_*i*_, and correspondingly, *x*_*i*_, which dictates the *k*-mers we should keep in our training data. Had we instead tried to use a random selection scheme, and post-hoc enforced coherence by removing *k*-mers that violated coherence, we would have ended up with dramatically fewer methylated sites in the training data (Supplementary Figure S2). Note that, for each value of *p, k*-mer-selection was performed only once across all 6 training folds within the 6-fold cross-validation used. Test sets were left as-is (with no *k*-mer-selection performed), other than as noted in the main paper.

Note that in a practical experiment intended to obtain limited training data for a caller, one might consider using this selection scheme to decide which portions of the genome to obtain labels for. However, our intent here was simply to simulate incomplete training data from our *k*-mer-complete 5mC data in order to investigate the effect of increasingly less complete data on different ML-based calling approaches. Our original intent had been to use a random selection scheme, but we then realized it was not coherent.

### Training with Nanopolish and DeepSignal

We used publicly available software for Nanopolish [10] with default settings. In this software, first the current time series in each Nanopore read is segmented into events, which are then aligned to a reference genome. Then, each *k*-mer is associated with a list of events. Second, these lists of events are then used to update the emission distribution parameters for each *k*-mer in the HMM. This process is repeated from the alignment step for five iterations.

We used software publicly provided by DeepSignal with default settings. For a given CpG site in the reference genome and a read covering the site, DeepSignal extracts a feature vector containing nucleotide sequence information, current summary statistics, and raw current values corresponding to a window centered on the CpG dinucleotide.

There are two notable side effects of our *k*-mer-selection downsampling. First, since we are only concerned with removing methylated sites from the data, our procedure naturally introduces significant class imbalance in the training data between methylated and unmethylated *k*-mers, since all unmethylated sites remained. Since the HMM is class-imbalance agnostic, we did not need to account for these issues for Nanopolish. However, for DeepSignal, we found it was needed. Consequently, to mitigate this bias, we correspondingly downsampled the negative (unmethylated) class at random, such that the proportion of each class was the same. Second, as we decreased *p*, the number of unique *k*-mers decreased, and correspondingly, the amount of training data. In order to keep the size of the training data (which can comprise multiple reads for the same *k*-mer), we always used the number of reads corresponding to the smallest value of *p* in our experiments, *p* = 5. Altogether, this left us for DeepSignal with around 1 million CpG sites for NA128, and 4 million for HX1 (which originally contained many more methylated sites than NA12878), for any value of *p*.

### Amortized-HMM

Our new approach, the Amortized-HMM, extends the HMM method for methylation calling through use of a DNN. From Nanpolish HMM training, we obtain emission distribution parameters (a scalar mean and a scalar variance) for *k*-mers observed at high enough frequency in the training data. We take these as labels for a feedforward deep neural network (FDNN) that learns a mapping from *k*-mers to emission distribution parameters. Next we impute any missing (*i. e*.,, those that Nanopolish left to the default) emission distributions with our FDNN, before using Nanpolish to make calls. In using only FDNN-based emission distributions by overriding all Nanopolish ones, performance was diminished (Supplementary Figure S3).

Our FDNN requires us to featurize each *k*-mer given as input to the FDNN, which we do as follows for each 6-mer. We use a one-hot encoding with the alphabet `{A, C, G, T, M}`, where `M` denotes a methylated cite, with the other letters denoting nucleotides, yielding 30 binary features. We also one-hot encode each dinucleotide formed from adjacent positions, yielding 105 binary features. Finally, we use a binary encoding of whether the position contains a C/M, or does not, yielding 6 more binary features. The motivation for including this last feature is that *M* is closely tied to *C* by way of being a modified cytosine, so we hypothesized that they may have similar effects on the nanopore current. In total, the feature vector was of length 141. Note that we additionally experimented with one-hot encoding trinucleotides from consecutive triplets in the *k*-mer, but this did not improve performance.

Rather than the more common losses used for training neural networks, we used a symmetrized Kullback-Leibler (KL) divergence loss to train our FDNN because the labels were themselves parameters of a distribution. Had we used say mean squared error, the numerical difference between the FDNN outputs and the labels would likely not meaningfully reflect the difference in distributions. In particular, we let *P* ∼ *N* (*µ, σ*^2^) be a Gaussian random variable with *µ* and *σ*^2^ estimated from Nanopolish (*i. e*., our supervisory labels), and let 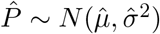 be a Gaussian variable with 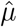, and 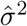 predicted by the Amortized-HMM. The symmetrized KL-divergence between these is defined as 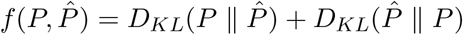, whose components can be computed in closed form by 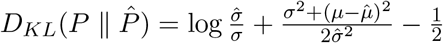 and 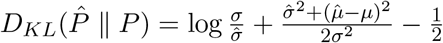.

Architecture search was performed for Amortized-HMM only for the *k*-mer complete setting, and then these parameters were used for all experiments, irrespective of the value of *p*. We determined the number of hidden layers, *d*, and the size of each hidden unit, *h*, with an 80%/20% random split of the data. We performed grid search over the hyperparameter values *d* ∈ {3, 4, 5, 6} and *h* ∈ {16, 32, 64, 128}.

### DeepSignal hyperparameter search for each *p*

For apples-to-apples comparison with Amortized-HMM, and to reduce the substantial computational burden, all reported experiments used architecture searches only from the *k*-mer-complete setting (*i. e*., that provided by DeepSignal), other than as noted next. As noted in the main text, we here performed an architecture search for DeepSignal, for a salient value of *p* = 10, to see if this would cause it to outperform Amortized-HMM(without doing this added architecture search for each *p*). The value *p* = 10 corresponds to a regime of low *k*-mer coverage and one where our approach substantially outperformed DeepSignal (Figure 1). We used the NA12878 dataset for training and validation. Furthermore, we tuned the same network hyperparameters that were originally tuned by Ni et al. [18]: the length of the *k*-mer context, the number of BRNN layers, and the number of inception layers. Despite doing so, DeepSignal did not outperform Amortized-HMM (Supplementary Figure S1).

## Acknowledgments

This research used the Savio computational cluster resource provided by the Berkeley Research Computing program at the University of California, Berkeley (supported by the UC Berkeley Chancellor, Vice Chancellor for Research, and Chief Information Officer). Partial support was provided by the Koret-UC Berkeley-Tel Aviv University Initiative in Computational Biology and Bioinformatics. Y.E. acknowledges the European Research Council consolidator grant (817811).

## Supplementary Material

### Supplementary Figures

**Figure S1:**
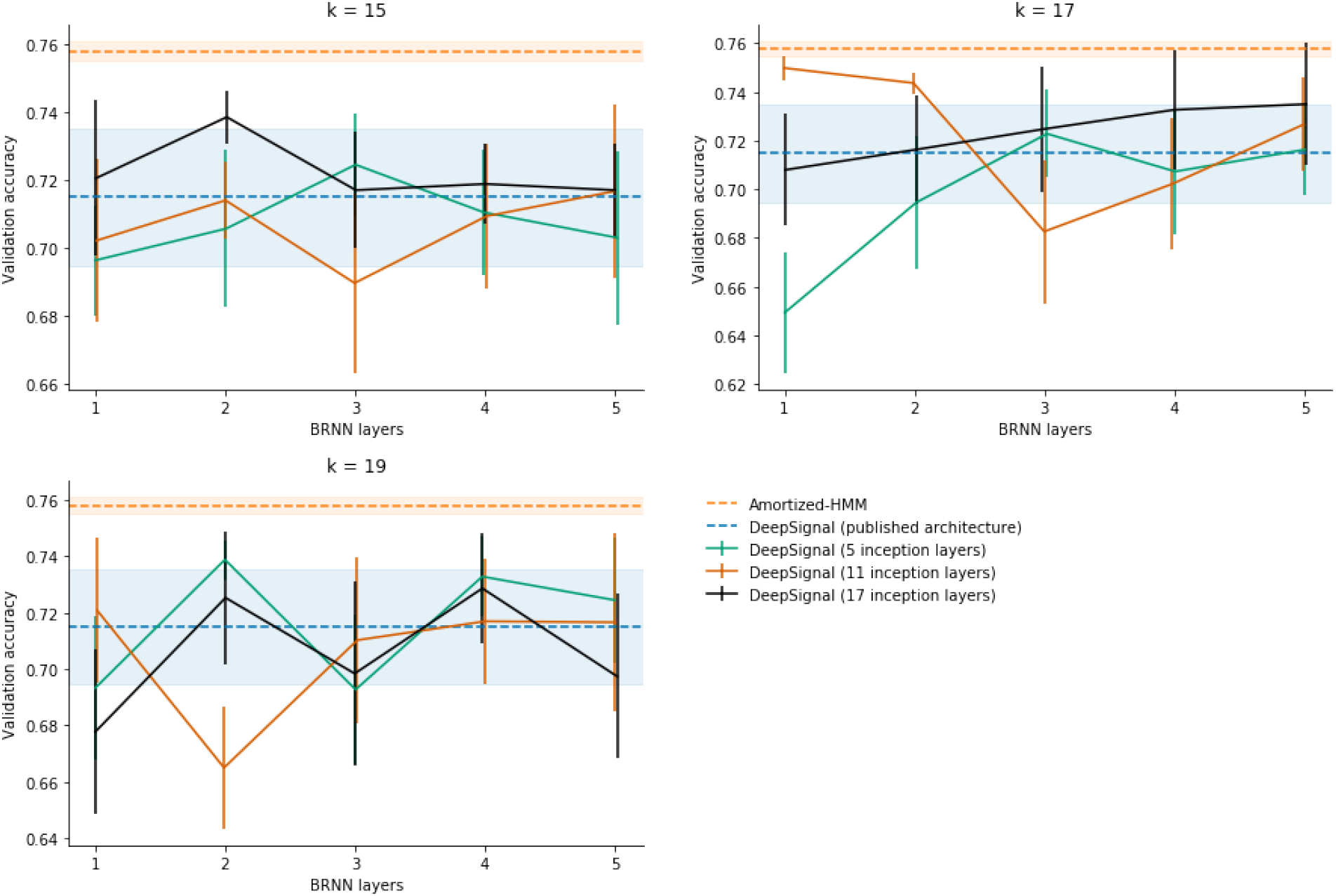
Architecture search for DeepSignal in *k*-mer-incomplete setting (*p* = 10). Results averaged over 6-fold cross-validation on the NA12878 dataset, with length of error bars and of the shaded regions equal to one standard deviation across the folds. The hyperparameter settings used in DeepSignal are 17 for the *k*-mer context length (*k*), 3 for the number of BRNN layers, and 11 for the number of inception layers. For improved readability, Gaussian noise was added onto the *x*-values for each data point.

**Figure S2:**
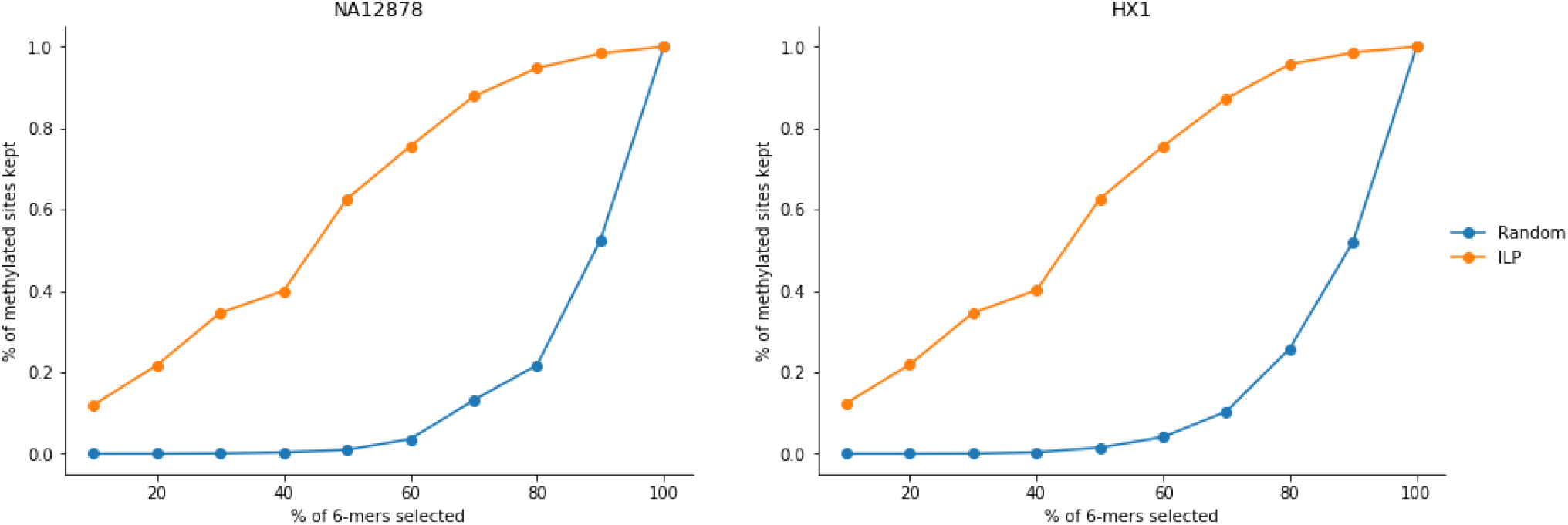
Comparison of random *k*-mer selection and ILP-based selection. Change in percentage of methylated sites retained in data with respect to percentage of *k*-mers kept, compared between randomly selected *k*-mer sets and ILP-selected *k*-mer sets.

**Figure S3:**
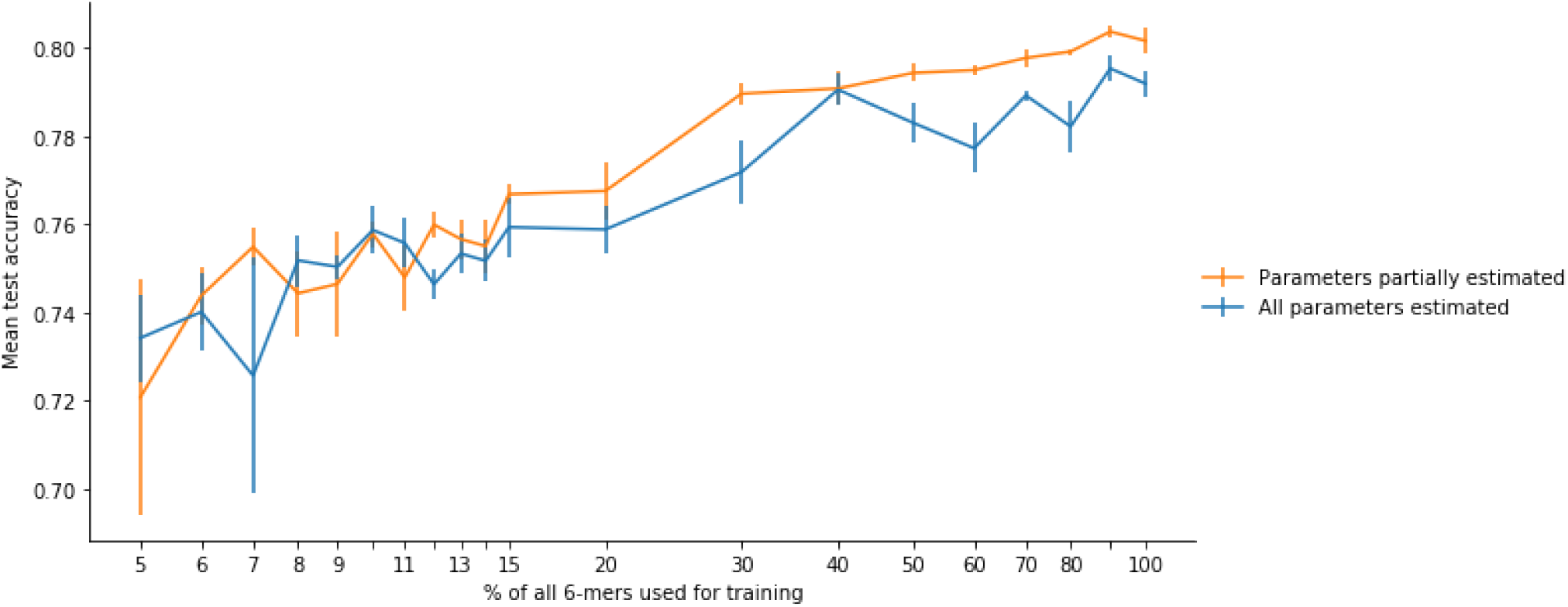
Accuracy of Amortized-HMM when all emission distribution parameters are estimated via FDNN versus when only parameters not updated during Nanopolish training phase are estimated. Results averaged over 6-fold cross-validation on the NA12878 dataset, with length of error bars equal to one standard deviation across the folds.

### Supplementary Tables

**Table S1:**
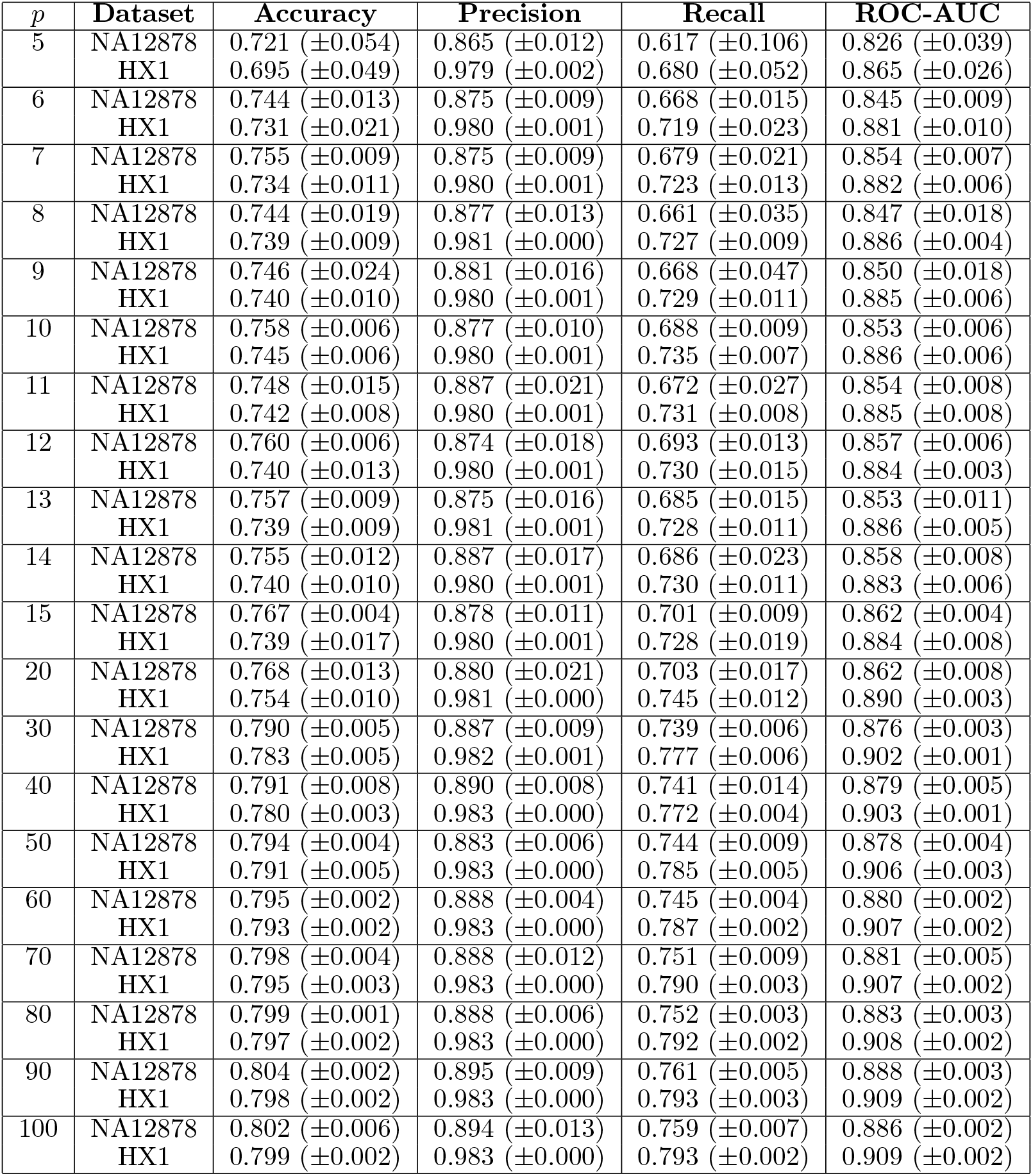
Performance metrics for Amortized-HMM for varying levels of *k*-mer incompleteness in the training data (*p*). Evaluations were performed on unfiltered, *k*-mer complete data. Mean and standard deviation values are reported from 6-fold cross-validation.

| $p$ | Dataset | Accuracy | Precision | Recall | ROC-AUC |
| --- | --- | --- | --- | --- | --- |
| 5 | NA12878 | 0.721 ($\pm 0.054$) | 0.865 ($\pm 0.012$) | 0.617 ($\pm 0.106$) | 0.826 ($\pm 0.039$) |
| | HX1 | 0.695 ($\pm 0.049$) | 0.979 ($\pm 0.002$) | 0.680 ($\pm 0.052$) | 0.865 ($\pm 0.026$) |
| 6 | NA12878 | 0.744 ($\pm 0.013$) | 0.875 ($\pm 0.009$) | 0.668 ($\pm 0.015$) | 0.845 ($\pm 0.009$) |
| | HX1 | 0.731 ($\pm 0.021$) | 0.980 ($\pm 0.001$) | 0.719 ($\pm 0.023$) | 0.881 ($\pm 0.010$) |
| 7 | NA12878 | 0.755 ($\pm 0.009$) | 0.875 ($\pm 0.009$) | 0.679 ($\pm 0.021$) | 0.854 ($\pm 0.007$) |
| | HX1 | 0.734 ($\pm 0.011$) | 0.980 ($\pm 0.001$) | 0.723 ($\pm 0.013$) | 0.882 ($\pm 0.006$) |
| 8 | NA12878 | 0.744 ($\pm 0.019$) | 0.877 ($\pm 0.013$) | 0.661 ($\pm 0.035$) | 0.847 ($\pm 0.018$) |
| | HX1 | 0.739 ($\pm 0.009$) | 0.981 ($\pm 0.000$) | 0.727 ($\pm 0.009$) | 0.886 ($\pm 0.004$) |
| 9 | NA12878 | 0.746 ($\pm 0.024$) | 0.881 ($\pm 0.016$) | 0.668 ($\pm 0.047$) | 0.850 ($\pm 0.018$) |
| | HX1 | 0.740 ($\pm 0.010$) | 0.980 ($\pm 0.001$) | 0.729 ($\pm 0.011$) | 0.885 ($\pm 0.006$) |
| 10 | NA12878 | 0.758 ($\pm 0.006$) | 0.877 ($\pm 0.010$) | 0.688 ($\pm 0.009$) | 0.853 ($\pm 0.006$) |
| | HX1 | 0.745 ($\pm 0.006$) | 0.980 ($\pm 0.001$) | 0.735 ($\pm 0.007$) | 0.886 ($\pm 0.006$) |
| 11 | NA12878 | 0.748 ($\pm 0.015$) | 0.887 ($\pm 0.021$) | 0.672 ($\pm 0.027$) | 0.854 ($\pm 0.008$) |
| | HX1 | 0.742 ($\pm 0.008$) | 0.980 ($\pm 0.001$) | 0.731 ($\pm 0.008$) | 0.885 ($\pm 0.008$) |
| 12 | NA12878 | 0.760 ($\pm 0.006$) | 0.874 ($\pm 0.018$) | 0.693 ($\pm 0.013$) | 0.857 ($\pm 0.006$) |
| | HX1 | 0.740 ($\pm 0.013$) | 0.980 ($\pm 0.001$) | 0.730 ($\pm 0.015$) | 0.884 ($\pm 0.003$) |
| 13 | NA12878 | 0.757 ($\pm 0.009$) | 0.875 ($\pm 0.016$) | 0.685 ($\pm 0.015$) | 0.853 ($\pm 0.011$) |
| | HX1 | 0.739 ($\pm 0.009$) | 0.981 ($\pm 0.001$) | 0.728 ($\pm 0.011$) | 0.886 ($\pm 0.005$) |
| 14 | NA12878 | 0.755 ($\pm 0.012$) | 0.887 ($\pm 0.017$) | 0.686 ($\pm 0.023$) | 0.858 ($\pm 0.008$) |
| | HX1 | 0.740 ($\pm 0.010$) | 0.980 ($\pm 0.001$) | 0.730 ($\pm 0.011$) | 0.883 ($\pm 0.006$) |
| 15 | NA12878 | 0.767 ($\pm 0.004$) | 0.878 ($\pm 0.011$) | 0.701 ($\pm 0.009$) | 0.862 ($\pm 0.004$) |
| | HX1 | 0.739 ($\pm 0.017$) | 0.980 ($\pm 0.001$) | 0.728 ($\pm 0.019$) | 0.884 ($\pm 0.008$) |
| 20 | NA12878 | 0.768 ($\pm 0.013$) | 0.880 ($\pm 0.021$) | 0.703 ($\pm 0.017$) | 0.862 ($\pm 0.008$) |
| | HX1 | 0.754 ($\pm 0.010$) | 0.981 ($\pm 0.000$) | 0.745 ($\pm 0.012$) | 0.890 ($\pm 0.003$) |
| 30 | NA12878 | 0.790 ($\pm 0.005$) | 0.887 ($\pm 0.009$) | 0.739 ($\pm 0.006$) | 0.876 ($\pm 0.003$) |
| | HX1 | 0.783 ($\pm 0.005$) | 0.982 ($\pm 0.001$) | 0.777 ($\pm 0.006$) | 0.902 ($\pm 0.001$) |
| 40 | NA12878 | 0.791 ($\pm 0.008$) | 0.890 ($\pm 0.008$) | 0.741 ($\pm 0.014$) | 0.879 ($\pm 0.005$) |
| | HX1 | 0.780 ($\pm 0.003$) | 0.983 ($\pm 0.000$) | 0.772 ($\pm 0.004$) | 0.903 ($\pm 0.001$) |
| 50 | NA12878 | 0.794 ($\pm 0.004$) | 0.883 ($\pm 0.006$) | 0.744 ($\pm 0.009$) | 0.878 ($\pm 0.004$) |
| | HX1 | 0.791 ($\pm 0.005$) | 0.983 ($\pm 0.000$) | 0.785 ($\pm 0.005$) | 0.906 ($\pm 0.003$) |
| 60 | NA12878 | 0.795 ($\pm 0.002$) | 0.888 ($\pm 0.004$) | 0.745 ($\pm 0.004$) | 0.880 ($\pm 0.002$) |
| | HX1 | 0.793 ($\pm 0.002$) | 0.983 ($\pm 0.000$) | 0.787 ($\pm 0.002$) | 0.907 ($\pm 0.002$) |
| 70 | NA12878 | 0.798 ($\pm 0.004$) | 0.888 ($\pm 0.012$) | 0.751 ($\pm 0.009$) | 0.881 ($\pm 0.005$) |
| | HX1 | 0.795 ($\pm 0.003$) | 0.983 ($\pm 0.000$) | 0.790 ($\pm 0.003$) | 0.907 ($\pm 0.002$) |
| 80 | NA12878 | 0.799 ($\pm 0.001$) | 0.888 ($\pm 0.006$) | 0.752 ($\pm 0.003$) | 0.883 ($\pm 0.003$) |
| | HX1 | 0.797 ($\pm 0.002$) | 0.983 ($\pm 0.000$) | 0.792 ($\pm 0.002$) | 0.908 ($\pm 0.002$) |
| 90 | NA12878 | 0.804 ($\pm 0.002$) | 0.895 ($\pm 0.009$) | 0.761 ($\pm 0.005$) | 0.888 ($\pm 0.003$) |
| | HX1 | 0.798 ($\pm 0.002$) | 0.983 ($\pm 0.000$) | 0.793 ($\pm 0.003$) | 0.909 ($\pm 0.002$) |
| 100 | NA12878 | 0.802 ($\pm 0.006$) | 0.894 ($\pm 0.013$) | 0.759 ($\pm 0.007$) | 0.886 ($\pm 0.002$) |
| | HX1 | 0.799 ($\pm 0.002$) | 0.983 ($\pm 0.000$) | 0.793 ($\pm 0.002$) | 0.909 ($\pm 0.002$) |

**Table S2:**
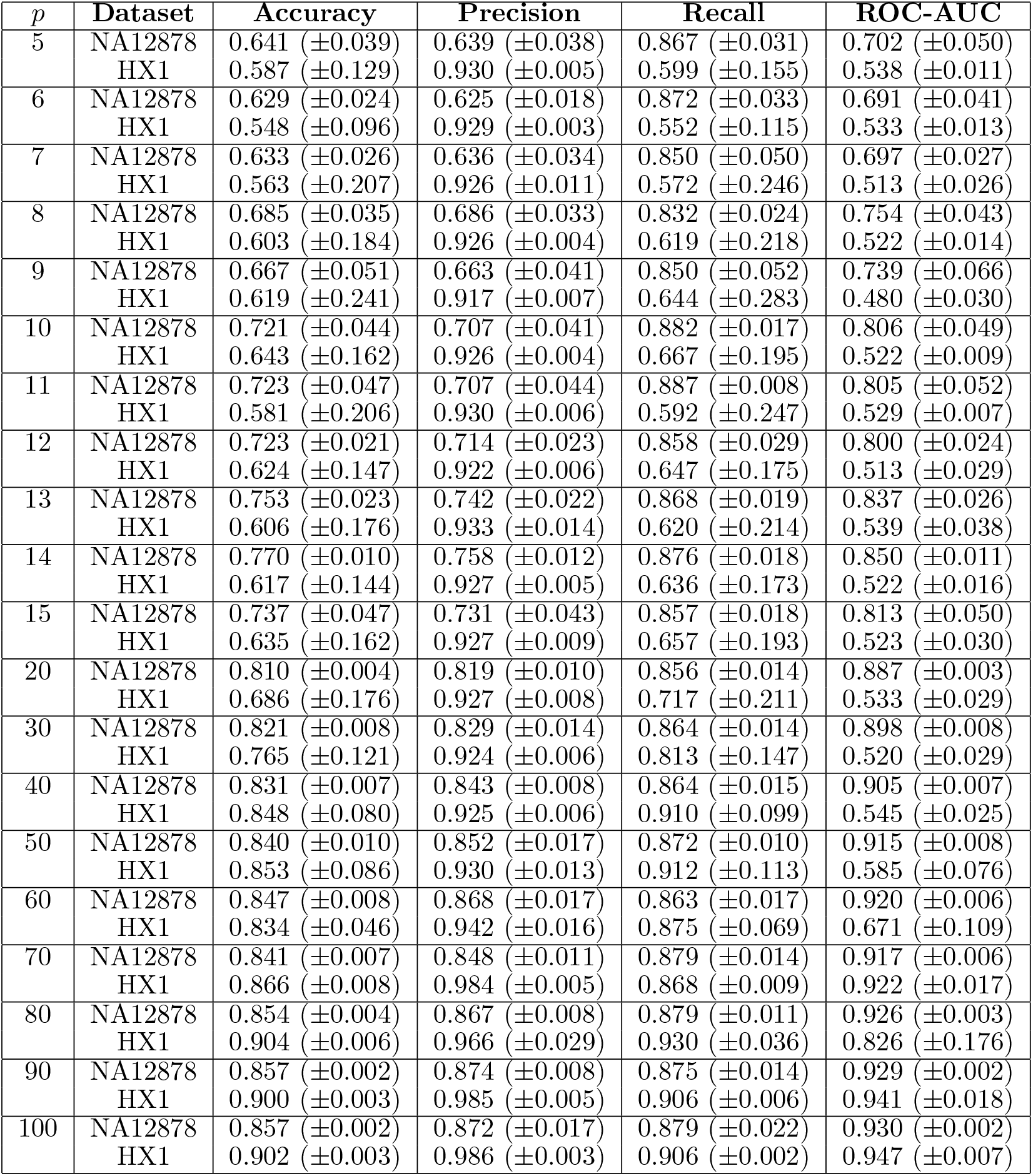
Performance metrics for DeepSignal for varying levels of *k*-mer incompleteness in the training data (*p*). Evaluations were performed on unfiltered, *k*-mer complete data. Mean and standard deviation values are reported from 6-fold cross-validation.

| $p$ | Dataset | Accuracy | Precision | Recall | ROC-AUC |
| --- | --- | --- | --- | --- | --- |
| 5 | NA12878 | 0.641 ($\pm 0.039$) | 0.639 ($\pm 0.038$) | 0.867 ($\pm 0.031$) | 0.702 ($\pm 0.050$) |
| | HX1 | 0.587 ($\pm 0.129$) | 0.930 ($\pm 0.005$) | 0.599 ($\pm 0.155$) | 0.538 ($\pm 0.011$) |
| 6 | NA12878 | 0.629 ($\pm 0.024$) | 0.625 ($\pm 0.018$) | 0.872 ($\pm 0.033$) | 0.691 ($\pm 0.041$) |
| | HX1 | 0.548 ($\pm 0.096$) | 0.929 ($\pm 0.003$) | 0.552 ($\pm 0.115$) | 0.533 ($\pm 0.013$) |
| 7 | NA12878 | 0.633 ($\pm 0.026$) | 0.636 ($\pm 0.034$) | 0.850 ($\pm 0.050$) | 0.697 ($\pm 0.027$) |
| | HX1 | 0.563 ($\pm 0.207$) | 0.926 ($\pm 0.011$) | 0.572 ($\pm 0.246$) | 0.513 ($\pm 0.026$) |
| 8 | NA12878 | 0.685 ($\pm 0.035$) | 0.686 ($\pm 0.033$) | 0.832 ($\pm 0.024$) | 0.754 ($\pm 0.043$) |
| | HX1 | 0.603 ($\pm 0.184$) | 0.926 ($\pm 0.004$) | 0.619 ($\pm 0.218$) | 0.522 ($\pm 0.014$) |
| 9 | NA12878 | 0.667 ($\pm 0.051$) | 0.663 ($\pm 0.041$) | 0.850 ($\pm 0.052$) | 0.739 ($\pm 0.066$) |
| | HX1 | 0.619 ($\pm 0.241$) | 0.917 ($\pm 0.007$) | 0.644 ($\pm 0.283$) | 0.480 ($\pm 0.030$) |
| 10 | NA12878 | 0.721 ($\pm 0.044$) | 0.707 ($\pm 0.041$) | 0.882 ($\pm 0.017$) | 0.806 ($\pm 0.049$) |
| | HX1 | 0.643 ($\pm 0.162$) | 0.926 ($\pm 0.004$) | 0.667 ($\pm 0.195$) | 0.522 ($\pm 0.009$) |
| 11 | NA12878 | 0.723 ($\pm 0.047$) | 0.707 ($\pm 0.044$) | 0.887 ($\pm 0.008$) | 0.805 ($\pm 0.052$) |
| | HX1 | 0.581 ($\pm 0.206$) | 0.930 ($\pm 0.006$) | 0.592 ($\pm 0.247$) | 0.529 ($\pm 0.007$) |
| 12 | NA12878 | 0.723 ($\pm 0.021$) | 0.714 ($\pm 0.023$) | 0.858 ($\pm 0.029$) | 0.800 ($\pm 0.024$) |
| | HX1 | 0.624 ($\pm 0.147$) | 0.922 ($\pm 0.006$) | 0.647 ($\pm 0.175$) | 0.513 ($\pm 0.029$) |
| 13 | NA12878 | 0.753 ($\pm 0.023$) | 0.742 ($\pm 0.022$) | 0.868 ($\pm 0.019$) | 0.837 ($\pm 0.026$) |
| | HX1 | 0.606 ($\pm 0.176$) | 0.933 ($\pm 0.014$) | 0.620 ($\pm 0.214$) | 0.539 ($\pm 0.038$) |
| 14 | NA12878 | 0.770 ($\pm 0.010$) | 0.758 ($\pm 0.012$) | 0.876 ($\pm 0.018$) | 0.850 ($\pm 0.011$) |
| | HX1 | 0.617 ($\pm 0.144$) | 0.927 ($\pm 0.005$) | 0.636 ($\pm 0.173$) | 0.522 ($\pm 0.016$) |
| 15 | NA12878 | 0.737 ($\pm 0.047$) | 0.731 ($\pm 0.043$) | 0.857 ($\pm 0.018$) | 0.813 ($\pm 0.050$) |
| | HX1 | 0.635 ($\pm 0.162$) | 0.927 ($\pm 0.009$) | 0.657 ($\pm 0.193$) | 0.523 ($\pm 0.030$) |
| 20 | NA12878 | 0.810 ($\pm 0.004$) | 0.819 ($\pm 0.010$) | 0.856 ($\pm 0.014$) | 0.887 ($\pm 0.003$) |
| | HX1 | 0.686 ($\pm 0.176$) | 0.927 ($\pm 0.008$) | 0.717 ($\pm 0.211$) | 0.533 ($\pm 0.029$) |
| 30 | NA12878 | 0.821 ($\pm 0.008$) | 0.829 ($\pm 0.014$) | 0.864 ($\pm 0.014$) | 0.898 ($\pm 0.008$) |
| | HX1 | 0.765 ($\pm 0.121$) | 0.924 ($\pm 0.006$) | 0.813 ($\pm 0.147$) | 0.520 ($\pm 0.029$) |
| 40 | NA12878 | 0.831 ($\pm 0.007$) | 0.843 ($\pm 0.008$) | 0.864 ($\pm 0.015$) | 0.905 ($\pm 0.007$) |
| | HX1 | 0.848 ($\pm 0.080$) | 0.925 ($\pm 0.006$) | 0.910 ($\pm 0.099$) | 0.545 ($\pm 0.025$) |
| 50 | NA12878 | 0.840 ($\pm 0.010$) | 0.852 ($\pm 0.017$) | 0.872 ($\pm 0.010$) | 0.915 ($\pm 0.008$) |
| | HX1 | 0.853 ($\pm 0.086$) | 0.930 ($\pm 0.013$) | 0.912 ($\pm 0.113$) | 0.585 ($\pm 0.076$) |
| 60 | NA12878 | 0.847 ($\pm 0.008$) | 0.868 ($\pm 0.017$) | 0.863 ($\pm 0.017$) | 0.920 ($\pm 0.006$) |
| | HX1 | 0.834 ($\pm 0.046$) | 0.942 ($\pm 0.016$) | 0.875 ($\pm 0.069$) | 0.671 ($\pm 0.109$) |
| 70 | NA12878 | 0.841 ($\pm 0.007$) | 0.848 ($\pm 0.011$) | 0.879 ($\pm 0.014$) | 0.917 ($\pm 0.006$) |
| | HX1 | 0.866 ($\pm 0.008$) | 0.984 ($\pm 0.005$) | 0.868 ($\pm 0.009$) | 0.922 ($\pm 0.017$) |
| 80 | NA12878 | 0.854 ($\pm 0.004$) | 0.867 ($\pm 0.008$) | 0.879 ($\pm 0.011$) | 0.926 ($\pm 0.003$) |
| | HX1 | 0.904 ($\pm 0.006$) | 0.966 ($\pm 0.029$) | 0.930 ($\pm 0.036$) | 0.826 ($\pm 0.176$) |
| 90 | NA12878 | 0.857 ($\pm 0.002$) | 0.874 ($\pm 0.008$) | 0.875 ($\pm 0.014$) | 0.929 ($\pm 0.002$) |
| | HX1 | 0.900 ($\pm 0.003$) | 0.985 ($\pm 0.005$) | 0.906 ($\pm 0.006$) | 0.941 ($\pm 0.018$) |
| 100 | NA12878 | 0.857 ($\pm 0.002$) | 0.872 ($\pm 0.017$) | 0.879 ($\pm 0.022$) | 0.930 ($\pm 0.002$) |
| | HX1 | 0.902 ($\pm 0.003$) | 0.986 ($\pm 0.003$) | 0.906 ($\pm 0.002$) | 0.947 ($\pm 0.007$) |

**Table S3:**
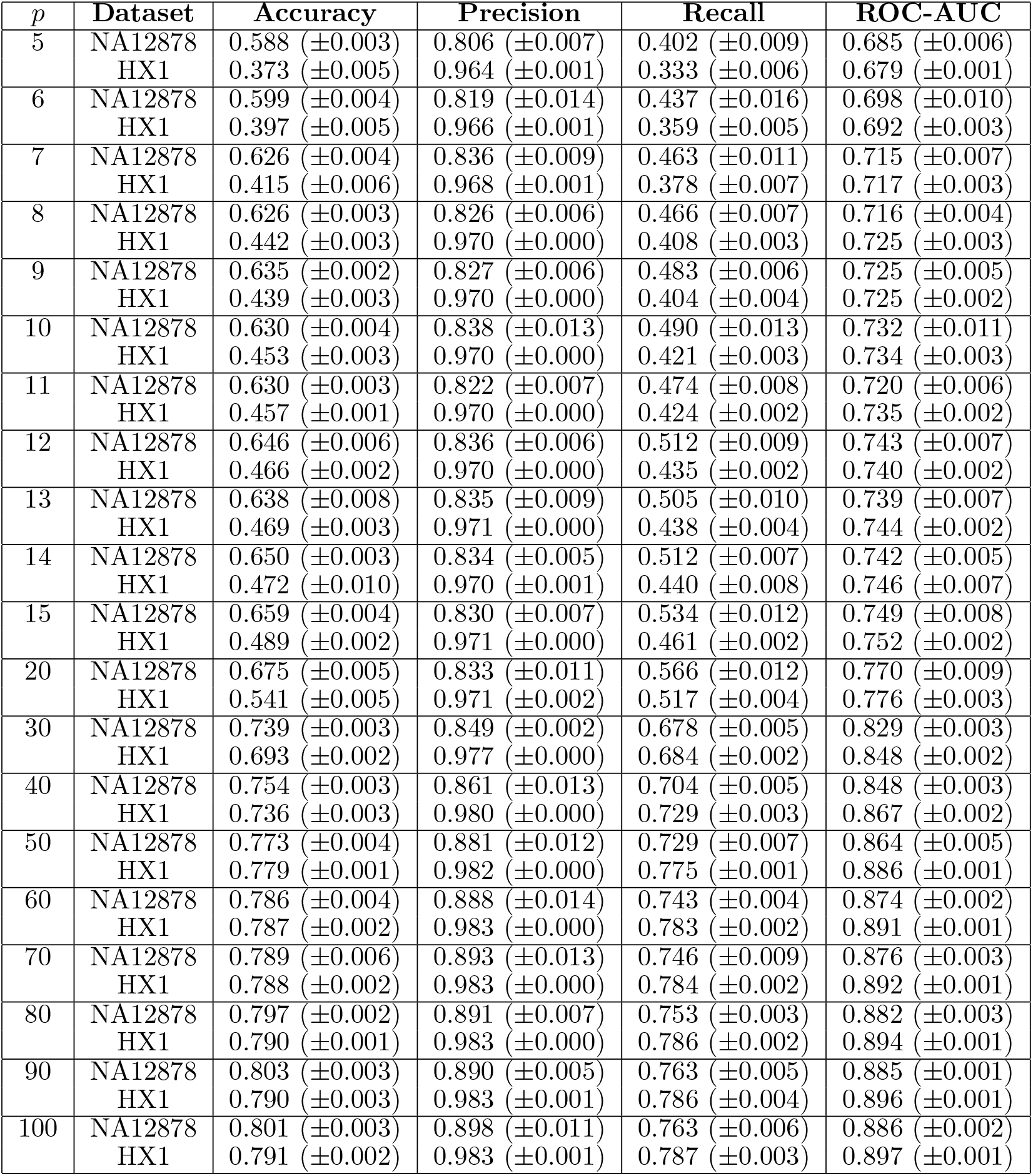
Performance metrics for Nanopolish for varying levels of *k*-mer incompleteness in the training data (*p*). Evaluations were performed on unfiltered, *k*-mer complete data. Mean and standard deviation values are reported from 6-fold cross-validation.

